# Tissue mechanics drives epithelialization, goblet cell regeneration, and restoration of a mucociliated epidermis on the surface of embryonic aggregates

**DOI:** 10.1101/696997

**Authors:** Hye Young Kim, Timothy R. Jackson, Carsten Stuckenholz, Lance A. Davidson

## Abstract

Injury, surgery, and disease often disrupt tissues and it is the process of regeneration that aids the restoration of architecture and function. Regeneration can occur through multiple strategies including induction of stem cell expansion, transdifferentiation, or proliferation of differentiated cells. We have uncovered a case of regeneration that restores a mucociliated epithelium from mesenchymal cells. Following disruption of embryonic tissue architecture and assembly of a compact mesenchymal aggregate, regeneration first involves restoration of an epithelium, transitioning from mesenchymal cells at the surface of the aggregate. Cells establish apico-basal polarity within 5 hours and a mucociliated epithelium within 24. Regeneration coincides with nuclear translocation of the putative mechanotransducer YAP1 and a sharp increase in aggregate stiffness, and regeneration can be controlled by altering stiffness. We propose that regeneration of a mucociliated epithelium occurs in response to biophysical cues sensed by newly exposed cells on the surface of a disrupted mesenchymal tissue.

## Main

Xenopus embryos develop a mucociliary epidermis that functions much like respiratory epithelia in mammals^1, 2^. Goblet cell progenitors develop exclusively from the epithelial surface layer whereas multiciliated and other accessory cells derive from deeper layer cells^3–6^ following notch-mediated patterning^7^. Once committed, multiciliated and accessory cell precursors intercalate between goblet cell progenitors to establish a fully functioning mucociliary epidermis^8^. Multiciliated and accessory cells have been extensively studied, yet the conditions driving goblet cell specification and the role of the mechanical microenvironment remain unclear.

Physical forces contribute to many cell fate decisions. For instance, during the first fate decision in the mouse embryo, polarized cell contractility generates asymmetric cell tension between superficial outer and inner deep cells leading to separation of trophectoderm and inner cell mass^9^. Physical forces or the deformations they generate also appear to pattern follicles in the skin^10^. *In vitro* studies suggest the tissue microenvironment, physically defined by factors such as stiffness, size, and substrate topology can direct stem cell lineage specification and renewal^11–13^; however, the contribution of such physicomechanical cues in embryonic cell specification is poorly understood. The gap between *in vivo* and *in vitro* studies reflects a paucity of model systems where tissue mechanics, cell behaviors, and cell fate choices can be studied quantitatively. To understand the role of tissue mechanics in embryonic cell specification, we investigated tissue mechanical properties during goblet cell regeneration on the surface of embryonic cell aggregates generated from early stage Xenopus laevis embryos.

Deep mesenchymal cells isolated from embryonic ectoderm and shaped into aggregates undergo an unexpected but profound transformation into an epithelial cell type. Embryonic cells isolated from deep layers of the Xenopus laevis embryo ectoderm, i.e. cells immediately below the simple epithelia of the ectoderm generate compact aggregates (Fig. 1a). Simple epithelia of the superficial cell layer assemble tight junctions^14^ and keratin intermediate filaments^15^ distinguishing them from deep mesenchymal cells and allowing the efficient separation of superficial layers from deep layer cells after brief exposure to calcium-magnesium-free media (Fig. 1a). Isolated deep ectoderm cells transferred to a non-adherent centrifuge tube rapidly adhere to each other in less than two hours to form a compact spherical aggregate. As expected, immunostaining of F-actin and fibronectin (FN) show regions where surface cells extend F-actin rich protrusions and assemble fibronectin fibrils (Fig. 1b, 1.5 hours post aggregation, hpa). However, by 5 hpa, clusters of cells on the aggregate surface are clear of FN and protrusions and adopt distinctive epithelial-like shapes with sharp cell boundaries marked by dense F-actin cables (Fig. 1b, see arrows). By 24 hpa, the entire surface develops into a mature epidermis, with multiciliated cells indicated by dense apical actin and a surface devoid of FN fibrils (Fig. 1b, Supplementary Fig. 1a). To rule out contamination by epithelial cells during microsurgery we surface labeled the outer cell layer of embryos used for making aggregates (Fig. 1c) and found no contaminating cells (Fig. 1d). Phenotypic transitions occurred across a range of aggregate sizes (Fig. 1e and f) from large (cells from 4 embryo-ectoderm explants) to small (cells from 1/4 of an ectoderm explant isolated from a single embryo). Thus, epithelial-like cells rapidly regenerate on the surface of a simple aggregate in the absence of externally provided factors.

**Figure 1:**
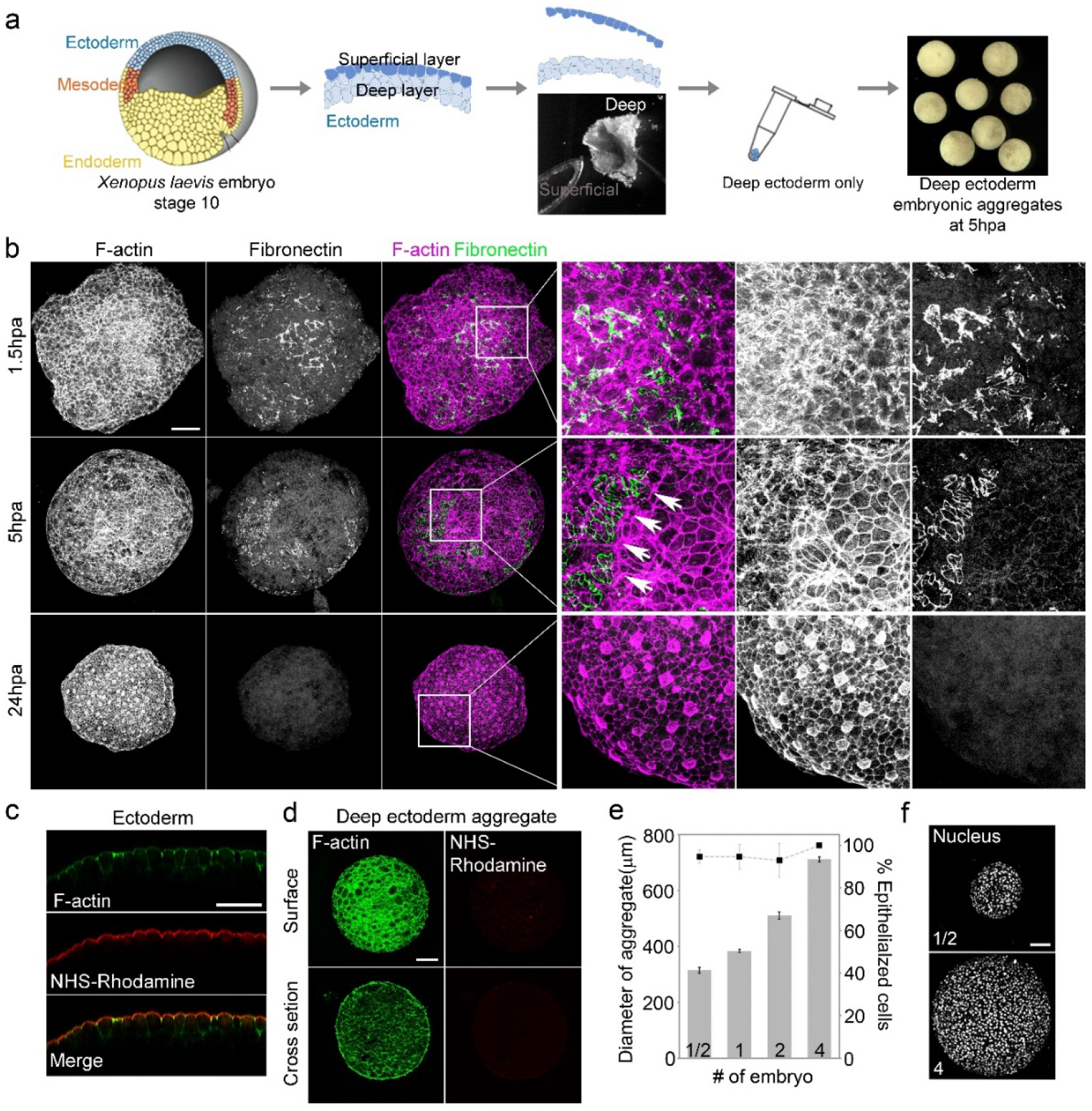
Surface cells of deep ectoderm aggregates undergo epithelial-like phenotypic transition. a, Schematic of the assembly of deep ectoderm cell aggregates from early Xenopus embryo (Stage 10). b, Surface F-actin and fibronectin (FN) from maximum intensity projections at 1.5, 5, and 24 hours post aggregation (hpa). Arrows indicate margin of FN where dense circumapical F-actin suggests epithelial cell phenotype. Scale bar, 100 μm. c, Transverse sectional view through the ectoderm of NHS-Rhodamine surface-labelled embryos. Scale bar, 50 μm. Rhodamine is restricted to the apical surface of outer epithelial cells. d, Deep ectoderm aggregates generated from NHS-Rhodamine surface-labelled embryos. Scale bar, 100 μm. Lack of rhodamine indicates absence of contaminating epithelia. e, Percent of epithelial cell phenotype found on the surface of different sized deep ectoderm aggregates at 24 hpa. Aggregates assembled with various number of embryo-ectoderm explants (1/2, 1, 2, and 4). f, Nuclei labeled in deep ectoderm aggregates from 1/2- and 4-embryo-ectoderm explant containing aggregates. Scale bar, 100 μm.

Apical-basal polarity is established progressively in cells exposed on the surface of the aggregate. Cell membranes facing the external media accumulate aPKC, an early marker of apical polarity by 5 hpa, continuing to 24 hpa when nuclei align in a pattern analogous to that seen in simple epithelia (Fig. 2a). By 5 hpa, epithelia-specific keratin intermediate filaments assemble along the apical surface; by 24 hpa, a mature epithelium forms with a dense keratin network (Fig. 2b). Epithelia-specific tight junction protein, ZO-1, also appears on the outer surface of a subpopulation of cells by 5 hpa and covers the entire surface by 24 hpa (Fig. 2c). Live imaging of GFP-ZO1 at early stages reveals scattered puncta as well as circumapical junctions (Fig. 2c and d) suggesting protrusive surface cells progressively transition to a tight epithelium. GFP-ZO-1 organizes on the surface of single cells and small groups as early as 2 hpa with the number of apically localized ZO-1 cells increasing over time, accounting for 10 to 15% of the surface area by 6 hpa (Fig. 2d and e).

**Figure 2:**
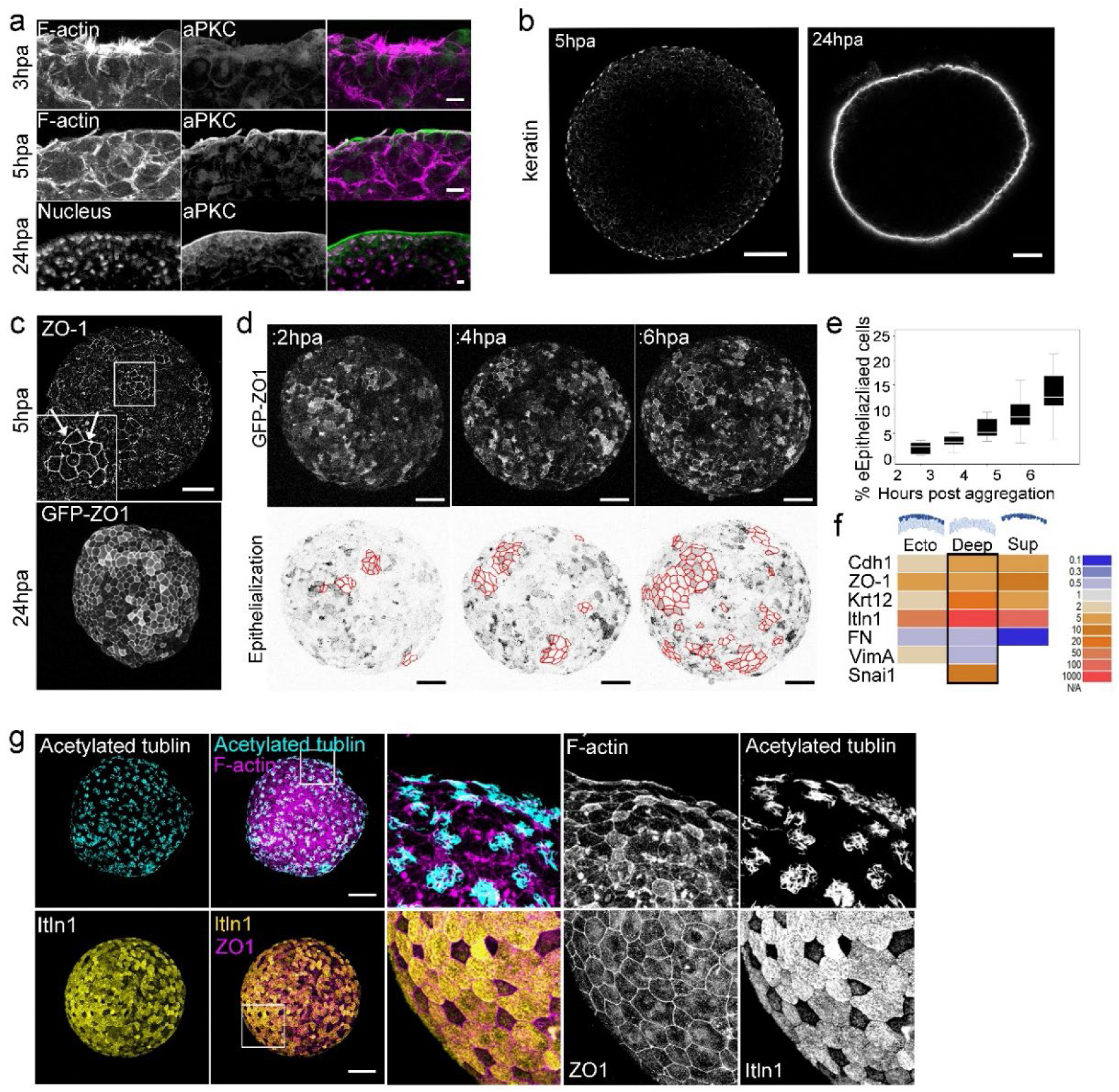
Epithelialization precedes differentiation of mucus-secreting goblet cells on the surface of deep ectoderm aggregates. a, Apical polarity protein aPKC localizes on the apical surface of aggregates by 5 hpa. Scale bar, 10 μm. b Apical localization of epithelial cytoskeleton keratin restricted to the outer surface of deep ectoderm aggregates. Cross sectional view of aggregates at 5 hpa (Scale bar, 100 μm) and 24 hpa (Scale bar, 50 μm). c, Maximum intensity projection of epithelial tight junctional protein ZO-1 expression in aggregates at 5 hpa (top; immunofluorescence staining) and 24 hpa (bottom; GFP-ZO-1 expression). Epithelialized cells marked by arrows (inset). Scale bar, 100 μm. d, Representative frames from a time-lapse sequence of an aggregate expressing GFP-ZO-1. Top panel: maximum intensity projection of GFP-ZO-1 shows cells undergo epithelialization (outlined with red on lookup-table-inverted images in lower panel) on the surface of aggregates from 2 to 6 hpa. Scale bar, 100 μm. e, The percent of cells having undergone epithelialization increases overtime (n=9 aggregates). f, qPCR expression profiling in deep ectoderm aggregates of epithelial (Cdh1, ZO-1, Krt12, and Itln 1) and mesenchymal (FN, VimA, and Snai1) genes for CT-based fold changes from 3 hpa to 24 hpa. g, At 24 hpa deep ectoderm aggregate is covered by epithelial cells including differentiated mucus secreting goblet cells (itln 1, Xeel) and radially intercalated multiciliated cells (acetylated tubulin). Scale bar, 100 μm.

The phenotypic transition to epithelial-like cells coincides with changing expression of epithelia-(Cdh1, ZO-1, Krt12. Itln1) and mesenchyme-specific genes (FN, VimA, and Snail) in ectoderm (both superficial and deep), deep ectoderm, and superficial ectoderm tissues (compare 3 and 24 hpa, Fig. 2f). Over this time-course, the aggregate increases expression of epithelial marker genes E-cadherin (Cdh1; 6-fold), ZO-1 (5-fold), and keratin (Krt12; (>25-fold) and the goblet cell marker intelectin 1 (Itln1; >1500-fold) while continuing to express mesenchymal genes. The continuing expression of mesenchymal genes in the aggregate reflects the continued presence of deep mesenchymal cells even as the surface epithelializes (Fig. 2f). Together these results indicate the surface of deep ectoderm aggregates regenerate an epithelial cell type.

How similar is this regenerated epithelial layer to the mature mucociliary epidermis found in the embryo? Xenopus larval epidermis forms as deep progenitors of multiciliated cells, small secretory cells, and ionocytes radially intercalate into the outer layer formed by a goblet cell precursors^6^. 24 hpa aggregates labeled with acetylated tubulin and F-actin reveal a pattern of multiciliated cells with dense apical actin cortex reminiscent of ciliated epithelium in similarly staged embryos (Fig. 2g). Furthermore, the ectoderm surface layer is dominated by mucus-secreting goblet cells marked by itln (Xeel, intelectin-1; Fig. 2g). We further ruled out a source of goblet cells from Notch-dependent fate decisions that generate accessory cell types *in vivo* (^7^; Supplementary Fig. 1). Thus, the newly epithelialized surface of aggregates regenerate goblet cells precursors that are fully competent to differentiate into patterned larval epidermis similar to that seen *in vivo.*

What triggers *de novo* epithelialization and goblet cell differentiation of the surface-layer cells in aggregates? Unlike deep ectoderm cells *in vivo,* deep ectoderm cells on the surface of aggregates are unconstrained by adjacent epithelia. Emergence of apparently random epithelial patches across the surface (Fig. 2c and d) suggests that cells respond to locally varying mechanical conditions. To test if cells might be responding to changes in their mechanical microenvironment, we analyzed the localization of Yes-associated protein 1 (YAP), a transcription factor whose nuclear translocation often correlates with changing mechanical conditions in other cell types, e.g. stress fiber formation^16^, cell shape change and ECM rigidity^17^, and stretch^18^. YAP is found at in the nucleus in cells on the surface of the aggregate (Fig. 3a) and depends on cell contractility (Fig. 3b). Nuclear levels of YAP increase in surface cells during the initial stages of epithelialization from 2 hpa to 5 hpa suggesting the surface cells experience higher tissue tension as they re-establish polarity (Fig. 3 c). Since reduced cell contractility reduced YAP nuclear translocation (Fig. 3b and c) we suspect that YAP reports an increase in tension at the aggregate surface.

**Figure 3:**
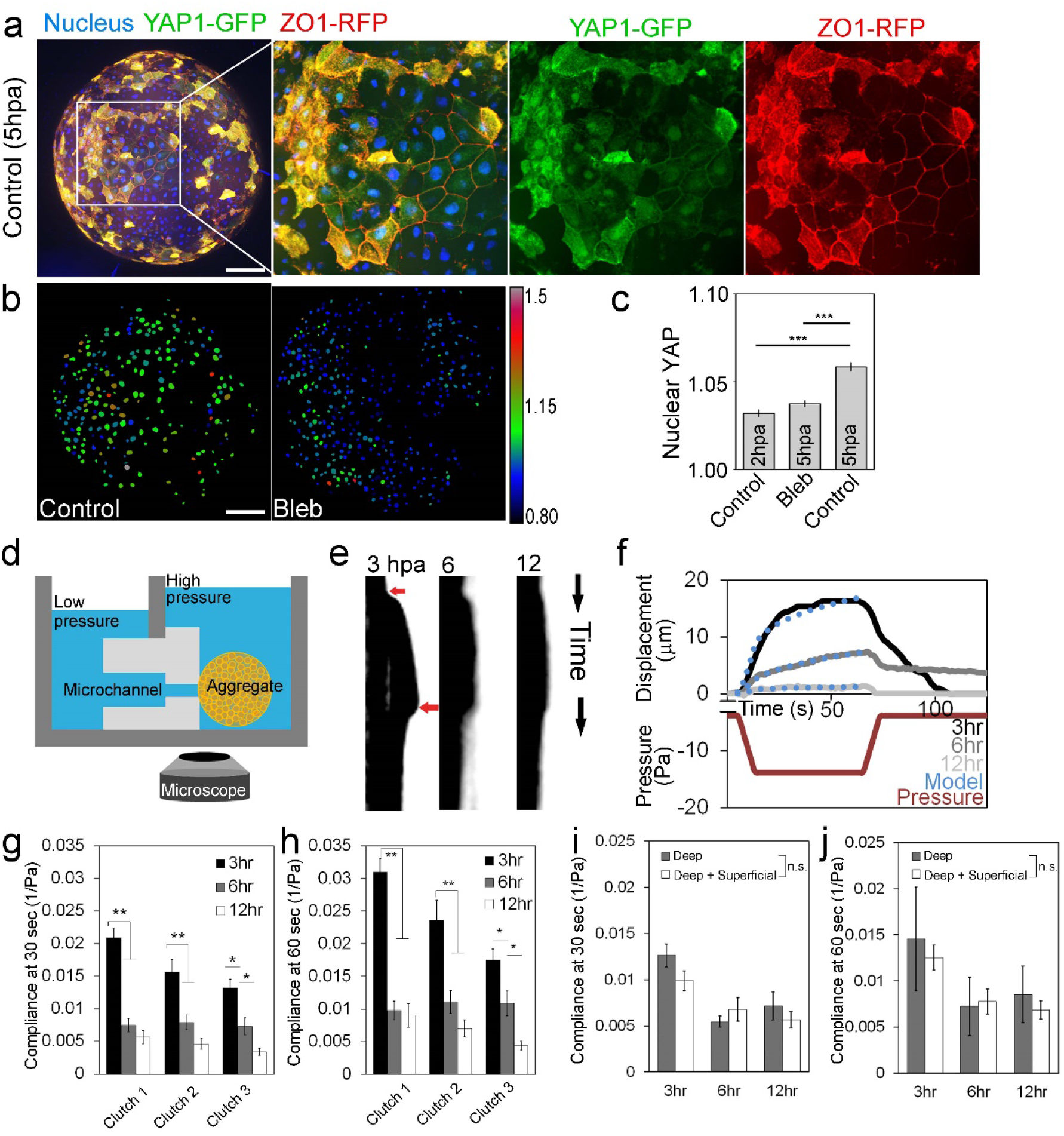
YAP nuclear translocation and tissue stiffening both coincide with epithelialization. a, Maximum intensity projection of deep ectoderm aggregate expressing YAP1-GFP and ZO-1-RFP and were stained for nucleus. Scale bar, 100 μm. b, Color coded YAP localization ratios for control and blebbistatin treated deep ectoderm aggregates at 5 hpa. Scale bar, 100 μm. c, YAP localization ratios of 5 hpa aggregates (n=2040 nuclei from 7 aggregates) are higher than blebbistatin treated aggregates (n=3471 nuclei from 7 aggregates) and 2 hpa aggregates (n=2167 nuclei from 7 aggregates). d, Schematic of micro-aspirator used to measure tissue compliance of aggregates by adjusting liquid pressure of the chamber through microchannel. e, Representative kymographs of tissue displacement over the length of the microaspiration experiments at 3, 6 and 12 hpa. Red arrows indicate when suction pressure is applied and then released. f, Representative graph of aspirated distance of an aggregate at 3 (black), 6 (dark gray), 12 hpa (light gray) relative to the pressure applied (red). The power-law model of creep compliance (blue dots) fit to this data. g-h, Creep compliance at 30 and 60 seconds indicating steady-state from micro-aspiration at 3 (black), 6 (gray) and 12 hpa (white). 11 to 16 aggregates were measured for each time point and repeated over three clutches (three clutches; * P<0.05, ** P<0.01, Error bar=+/− SE). i-j, Creep compliance at 30 and 60 seconds from micro-aspiration of aggregates consisting of only mesenchymal cells (gray) or both mesenchymal and superficial epithelial cells (white) 3, 6, and 12 hours post aggregation (one clutch, n=7 to 8 aggregates each). Differences in compliance are not statistically significant (Mann-Whitney U-test).

To quantify mechanical changes in the microenvironment of deep ectoderm aggregates, we measured the creep compliance, a measure of tension and tissue stiffness, using microaspiration^19^. Application of a small negative pressure (−10.1 Pa) to a small patch on the surface of the aggregate reports compliance to a depth of ~ 125 μm (Fig. 3d-f). Deep ectoderm aggregates at 3, 6, and 12 hpa, respectively, before, during, and after onset of epithelialization exhibit significant and large decreases in compliance, i.e. increased stiffness, over the first 6 hours, coincident with epithelization; compliance continues to decrease but appears to stabilize by later phases of differentiation (6 to 12 hpa, Fig. 3g and h). Comparing compliance of early aggregates (3 hpa) with or without superficial epithelia indicates that stiffness increases are not merely a consequence of epithelialization (Fig. 3i and j). Nuclear translocation of YAP together with phased reductions in compliance of the aggregates suggests that changes in the mechanical microenvironment promote surface epithelization in aggregates.

To understand how mechanics might control epithelialization and goblet cell differentiation we sought to test the roles of actomyosin contractility and cell-cell adhesion, key mediators of tissue mechanics in embryos^20^. To reduce contractility we expanded our earlier perturbations of cell contractility by incubating aggregates in either a Myosin II inhibitor, blebbistatin (100 μM) or a Rho-Kinase inhibitor, Y27632 (50 μM). To reduce cell-cell adhesion we expressed mutant forms of C-cadherin (cdh3), the major cadherin expressed within deep ectoderm cells at these stages, which lacked either the cytoplasmic (ΔC-C-cadherin) or extracellular domains (ΔE-C-cadherin)^21^. By 5 hpa approximately 10% of the surface area of untreated aggregates adopt an epithelial phenotype (Fig. 4a and b) as quantified from either F-actin or ZO-1 labeling where epithelial cells connected by tight junctions are enriched with circumapical F-actin along the boundary of the cells (Supplementary Fig. 2). Globally inhibiting contractility strongly reduces the epithelialized area leaving the surface covered by F-actin rich protrusions (Fig. 4a) like that seen at earlier stages (compare to Fig. 1b). To confirm the specific effects of the small molecule inhibitors of contractility, we expressed a mutant myosin binding subunit (MBS, MYPT1, or formally PPP1R12A), MBS^T695A^, which is known to inhibit contractility and subsequently increase tissue compliance^22^. Expression of MBS^T695A^ blocked epithelialization (Fig. 4a and b). Reducing cell-cell adhesion by moderate overexpression of ΔE-C-cadherin also blocked epithelialization (Fig. 4a and b) as did moderate overexpression of ΔC-C-cadherin (Fig. 4a and b, ΔC-C-cadherin Myc positive); we note adjacent wild-type cells (ΔC-C-cadherin, Myc negative) retain the ability to transition to and become epithelial (Supplementary Fig. 4). The reduced incidence of surface epithelialization following reductions in contractility or adhesion suggests both are required to advance epithelization in the aggregates.

**Figure 4:**
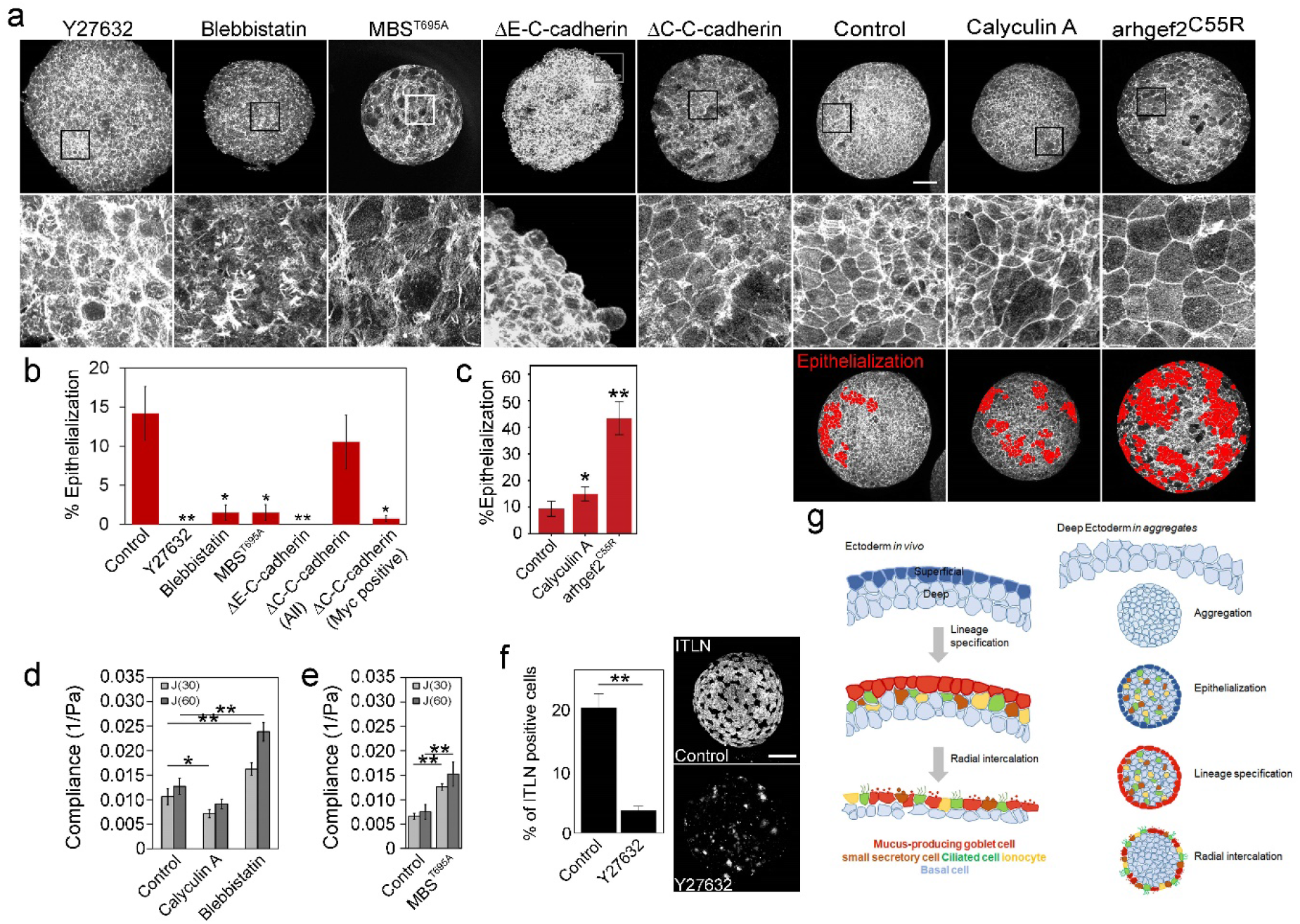
Contractility regulates surface epithelialization and goblet cell specification in deep ectoderm aggregates. a, Maximum intensity projection of F-actin stained aggregates at 5 hpa (control, n=30). Insets shown in lower panels. Scale bar, 100 μm. b, Epithelialization is reduced after lowering contractility (Y27632, n=9; Blebbistatin, n=10; MBS^T695A^; n=12) and altering cell-cell adhesion (ΔE-C-cadherin, n=15 and ΔC-C-cadherin, n=22). Analysis for mosaic ΔC-C-cadherin expression (anti-Myc positive) are shown in separate bar. (See Supplemental Experimental Procedure for analysis details). c, Epithelialization is increased after increasing contractility (Calyculin A, n=12; arhgef2^C55R^, n=10). Red filled cells indicate epithelialized cells and their surface area are quantified in a bar graph. Significance of each treatment from the control is shown by the asterisk (no asterisk, P>0.05; *, P<0.05; **, P<0.01; Error bar=+/− SE). d, Creep compliance at 30 and 60 seconds by micro-aspiration at 6 hpa after 4 hours of small molecule inhibitor treatment. Data represents four clutches pooled with 22 to 25 aggregates for each treatment. e, Creep compliance for MBS^T695A^ expressing aggregates. Data represents 8 aggregates (one clutch) per treatment. f, Percent of itln 1 positive goblet cells at 24 hpa in deep ectoderm aggregates (control, n=8; Y27632, n=10). Scale bar, 100 μm. g, Schematic contrasting developmental sequence of native embryonic ectoderm with *in vitro* regeneration of surface goblet cells in deep ectoderm aggregates. Regenerated epithelium serves as a substrate for radial intercalation of multiple cell types including multiciliated cells, ionocytes, and small secretory cells. Regenerated epithelial cells differentiate into mucus secreting goblet cells.

Since reduced contractility inhibited epithelialization, we wondered whether increasing contractility might accelerate epithelialization. To increase contractility we incubated aggregates with Calyculin A (10 nM), a myosin light chain phosphatase inhibitor known to increase contractility^23, 24^ and also expressed a potent activator of myosin contractility, a constitutively active mutant form of arhgef2 (arhgef2^C55R^), a RhoA-specific guanine nucleotide exchange factor known to strongly lower tissue compliance^25^. Both treatments sharply increased epithelialization (1.6-fold by Calyculin A and 4.6-fold by arhgef2^C55R^; Fig. 4a and c). The effects of arhgef2^C55R^ can be attributed to cell contractility, since its effects on epithelialization can be completely abolished by incubation with Y27632 (Supplementary Fig. S3a and b). Microaspiration further confirms that Calyculin A reduces compliance (Fig. 4d) and both Blebbistatin (Fig. 4d) and MBS^T695A^ (Fig. 4e) increase tissue compliance compared to control aggregates in concordance with their effects on actomyosin contractility. Furthermore, reduction of actomyosin contractility with Y27632 blocks goblet cell differentiation (Fig. 4f) indicating that tissue mechanics plays a role in both epithelization and differentiation; however it remains unclear whether epithelialization is a prerequisite for goblet cell differentiation. In summary, our data shows that contractility and tissue compliance regulate the onset of epithelialization and regeneration of a mucociliary epithelia in embryonic aggregates (Fig. 4g).

Mesenchymal Xenopus embryonic deep ectodermal cell aggregates regenerate a superficial epithelial layer of goblet cell precursors in as little as 5 hours. The phenotypic transition occurs in the absence of externally provided factors and is independent of endogenous patterning processes such as Notch-pathways that normally generate accessory cells in the deep ectoderm^26^. Unlike progenitor cells within stratified epithelia or pseudostratified epithelia^27^, native deep ectoderm cells remain in the deep layer through tadpole stages^28^. By contrast, endogenous goblet cell precursors originate from and are retained in the superficial layer of the ectoderm^6^. Thus, the programs driving epithelialization and regeneration of goblet cell precursors appear to be distinct from endogenous developmental programs.

We are surprised that the genetic networks regulating goblet cell differentiation are distinct from the pathways generating multiciliated and accessory cells^29^. Our study suggests a novel goblet cell differentiation pathway is notch-independent and regulated by the mechanical microenvironment. The sensitivity of precursor cells to mechanical cues may be responsible for pathological cases of basal cell differentiation where mucus secreting goblet cells are over- or under-produced^30^.

The mechanical microenvironment surrounding a progenitor cell can play a major role in determining the differentiation potential^11, 31^ and maintenance of stem cell populations^31^. What mechanical cues drive epithelialization and regeneration of goblet cells? Formation of an aggregate involves cell-cell contact, increasing cell-cell adhesion, and cell-autonomous actomyosin contractility. While all cells generate cortical contractions and membrane extensions into their surroundings, cells positioned within aggregates are bounded on all sides by cell-cell adhesions, whereas cells positioned along the surface of aggregates have two faces: one that is bounded by cell-cell adhesion and one that unbounded and open to culture media. As the mass compacts, cell projections toward the open surface would move laterally to contact neighboring cells in a manner similar to filopodial extensions during compaction early in mouse embryogenesis^32, 33^. We propose that as cell aggregates become tightly adhered they decrease in tissue compliance (Fig. 3d-j) and trigger the establishment of a novel apical-basal axis in surface cells which subsequently assemble epithelial specific junctions. The progressive polarization and assembly of apical junctions parallels critical events in the earliest stages of mammalian differentiation as the inner cell mass segregates into epithelial and deeper mesenchymal cell layers leading to the first cell steps of fate specification^34^. Based on our findings we speculate that strong apical-basal cues might trigger YAP localization but cannot rule out the possibility that polarization is a consequence of YAP nuclear translocation.

Xenopus ectoderm has been used historically to dissect fundamental signaling and patterning networks of development and stem cell differentiation^35, 36^. Biomechanical studies of Xenopus ectoderm have demonstrated how mechanics plays a direct role in regulating intercalation^37^, cell division^38^, epiboly^39^, and directional beating of multiciliated cells^40^. Mechanical and biophysical approaches in Xenopus will allow the field to revisit critical questions of patterning and induction to understand how mechanics integrates with canonical patterning systems.

## Acknowledgements

We thank Joe Shawky, Deepthi Vijayraghavan, Holley Lynch, Takehiko Ichikawa and other members of the group for their comments and discussions. This work was supported by the National Science Foundation (CBET-1547790) and the National Institutes of Health (R01HD044750; R56HL13495). TRJ was supported in part by the Cardiovascular Bioengineering Training Program (NIH NHLBI T32 HL076124). Any opinions, findings, and conclusions or recommendations expressed in this material are those of the authors and do not necessarily reflect the views of the National Science Foundation or the National Institutes of Health. We thank the Developmental Studies Hybridoma Bank at the University of Iowa, supported by the NICHD of the NIH, for the anti-keratin antibody 1H5.

## References

1. Walentek, P. & Quigley, I.K. What we can learn from a tadpole about ciliopathies and airway diseases: using systems biology in Xenopus to study cilia and mucociliary epithelia. genesis 55, e23001 (2017).

2. Dubaissi, E. & Papalopulu, N. Embryonic frog epidermis: a model for the study of cell-cell interactions in the development of mucociliary disease. Disease models and mechanisms 4, 179–192 (2011).

3. Quigley, I.K., Stubbs, J.L. & Kintner, C. Specification of ion transport cells in the Xenopus larval skin. Development 138, 705–714 (2011).

4. Stubbs, J., Vladar, E., Axelrod, J. & Kintner, C. Multicilin promotes centriole assembly and ciliogenesis during multiciliate cell differentiation. Nature cell biology 14, 140–147 (2012).

5. Walentek, P. et al. A novel serotonin-secreting cell type regulates ciliary motility in the mucociliary epidermis of Xenopus tadpoles. Development 141, 1526–1533 (2014).

6. Dubaissi, E. et al. A secretory cell type develops alongside multiciliated cells, ionocytes and goblet cells, and provides a protective, anti-infective function in the frog embryonic mucociliary epidermis. Development 141, 1514–1525 (2014).

7. Deblandre, G.A., Wettstein, D.A., Koyano-Nakagawa, N. & Kintner, C. A two-step mechanism generates the spacing pattern of the ciliated cells in the skin of Xenopus embryos. Development 126, 4715–4728 (1999).

8. Stubbs, J.L., Davidson, L., Keller, R. & Kintner, C. Radial intercalation of ciliated cells during Xenopus skin development. Development 133, 2507–2515 (2006).

9. Maître, J.-L. et al. Asymmetric division of contractile domains couples cell positioning and fate specification. Nature 536, 344 (2016).

10. Shyer, A.E. et al. Emergent cellular self-organization and mechanosensation initiate follicle pattern in the avian skin. Science 357, 811–815 (2017).

11. Engler, A.J., Sen, S., Sweeney, H.L. & Discher, D.E. Matrix elasticity directs stem cell lineage specification. Cell 126, 677–689 (2006).

12. McBeath, R., Pirone, D.M., Nelson, C.M., Bhadriraju, K. & Chen, C.S. Cell shape, cytoskeletal tension, and RhoA regulate stem cell lineage commitment. Dev Cell 6, 483–495 (2004).

13. Kilian, K.A., Bugarija, B., Lahn, B.T. & Mrksich, M. Geometric cues for directing the differentiation of mesenchymal stem cells. Proc Natl Acad Sci U S A 107, 4872–4877 (2010).

14. Merzdorf, C.S., Chen, Y.H. & Goodenough, D.A. Formation of functional tight junctions in Xenopus embryos. Dev Biol 195, 187–203 (1998).

15. Jamrich, M., Sargent, T.D. & Dawid, I.B. Cell-type-specific expression of epidermal cytokeratin genes during gastrulation of *Xenopus laevis*. Genes Dev 1, 124–132 (1987).

16. Chanet, S. & Martin, A.C. Mechanical force sensing in tissues. Progress in molecular biology and translational science 126, 317 (2014).

17. Dupont, S. et al. Role of YAP/TAZ in mechanotransduction. Nature 474, 179–183 (2011).

18. Aragona, M. et al. A mechanical checkpoint controls multicellular growth through YAP/TAZ regulation by actin-processing factors. Cell 154, 1047–1059 (2013).

19. von Dassow, M., Strother, J.A. & Davidson, L.A. Surprisingly simple mechanical behavior of a complex embryonic tissue. PLoS One 5, e15359 (2010).

20. Turlier, H. & Maître, J.-L. Mechanics of tissue compaction. Seminars in cell & developmental biology 47–48, 110-117 (2015).

21. Kurth, T. et al. Immunocytochemical studies of the interactions of cadherins and catenins in the early Xenopus embryo. Dev Dyn 215, 155–169 (1999).

22. Jackson, T.R., Kim, H.Y., Balakrishnan, U.L., Stuckenholz, C. & Davidson, L.A. Spatiotemporally controlled mechanical cues drive progenitor mesenchymal-to-epithelial transition enabling proper heart formation and function. Curr Biol (2017).

23. Yam, P.T. et al. Actin-myosin network reorganization breaks symmetry at the cell rear to spontaneously initiate polarized cell motility. J Cell Biol 178, 1207–1221 (2007).

24. Kim, H.Y. & Davidson, L.A. Punctuated actin contractions during convergent extension and their permissive regulation by the non-canonical Wnt-signaling pathway. Journal of Cell Science 124, 635–646 (2011).

25. Zhou, J., Kim, H.Y., Wang, J.H.-C. & Davidson, L.A. Macroscopic stiffening of embryonic tissues via microtubules, Rho-GEF, and assembly of contractile bundles of actomyosin. Development 137, 2785–2794 (2010).

26. Werner, M. & Mitchell, B. Understanding ciliated epithelia: the power of Xenopus. Genesis 50, 176–185 (2012).

27. Bragulla, H.H. & Homberger, D.G. Structure and functions of keratin proteins in simple, stratified, keratinized and cornified epithelia. Journal of anatomy 214, 516–559 (2009).

28. Schroeder, T.E. Neurulation in Xenopus laevis. An analysis and model based upon light and electron microscopy. J. Embryol. and Expt. Morphol. 23(2), 427–462 (1970).

29. Whitsett, J.A. Airway epithelial differentiation and mucociliary clearance. Annals of the American Thoracic Society 15, S143–S148 (2018).

30. Rock, J.R., Randell, S.H. & Hogan, B.L. Airway basal stem cells: a perspective on their roles in epithelial homeostasis and remodeling. Disease models & mechanisms 3, 545–556 (2010).

31. Gilbert, P.M. et al. Substrate elasticity regulates skeletal muscle stem cell self-renewal in culture. Science 329, 1078–1081 (2010).

32. Fierro-Gonzalez, J.C., White, M.D., Silva, J.C. & Plachta, N. Cadherin-dependent filopodia control preimplantation embryo compaction. Nat Cell Biol 15, 1424–1433 (2013).

33. Maître, J.-L., Niwayama, R., Turlier, H., Nédélec, F. & Hiiragi, T. Pulsatile cell-autonomous contractility drives compaction in the mouse embryo. Nature cell biology (2015).

34. Korotkevich, E. et al. The apical domain is required and sufficient for the first lineage segregation in the mouse embryo. Developmental cell 40, 235–247. e237 (2017).

35. Kuroda, H., Fuentealba, L., Ikeda, A., Reversade, B. & De Robertis, E.M. Default neural induction: neuralization of dissociated Xenopus cells is mediated by Ras/MAPK activation. Genes & development 19, 1022–1027 (2005).

36. Ariizumi, T. & Asashima, M. In vitro induction systems for analyses of amphibian organogenesis and body patterning. Int J Dev Biol 45, 273–279 (2001).

37. Sedzinski, J., Hannezo, E., Tu, F., Biro, M. & Wallingford, J.B. Emergence of an Apical Epithelial Cell Surface In Vivo. Developmental cell 36, 24–35 (2016).

38. Stooke-Vaughan, G.A., Davidson, L.A. & Woolner, S. Xenopus as a model for studies in mechanical stress and cell division. Genesis 55, e23004 (2017).

39. Stepien, T., Lynch, H.E., Yancey, S.X., Dempsey, L. & Davidson, L.A. Using a continuum model to decipher the mechanics of embryonic tissue spreading from time-lapse image sequences: An approximate Bayesian computation approach. PLoS One, 460774 (In Press).

40. Chien, Y.-H., Keller, R., Kintner, C. & Shook, D.R. Mechanical Strain Determines the Axis of Planar Polarity in Ciliated Epithelia. Current Biology 25, 2774–2784 (2015).

