## Supplementary Figures and Methods for "Tissue mechanics drives epithelialization, goblet cell regeneration, and restoration of a mucociliated epidermis on the surface of embryonic aggregates"

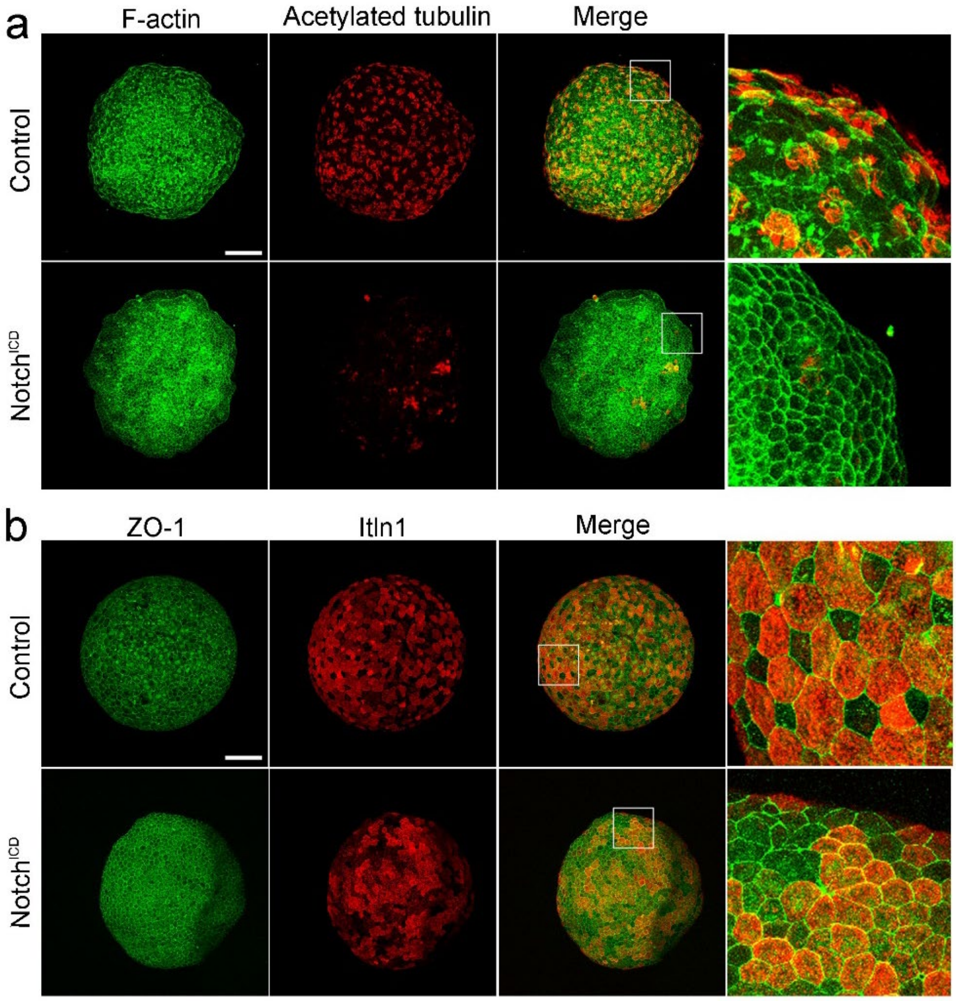


**Supplementary Figure 1: Epithelialization and goblet cell specification is independent from Notch signaling.**

a-b, Activation of Notch signaling by over-expression of the intracellular domain of $Notch^{ICD}$ inhibits the differentiation of multiciliated cells but does not inhibit regeneration of goblet cells in deep ectoderm aggregates (24 hpa). Scale bars, 100 µm.

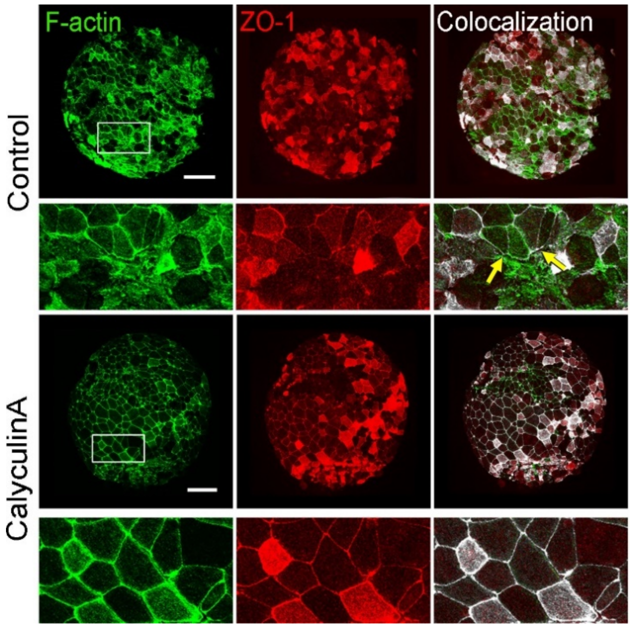


**Supplementary Figure 2: F-actin and ZO-1 co-localize on the boundary of epithelial cells.** Aggregates expressing ZO-1 RFP (red) and stained for F-actin (green) reveal co-localization of tight junctions and circumapical actin in epithelialized cells (white) in both control and Calyculin A treated aggregates. Scale bar, 100 μm.

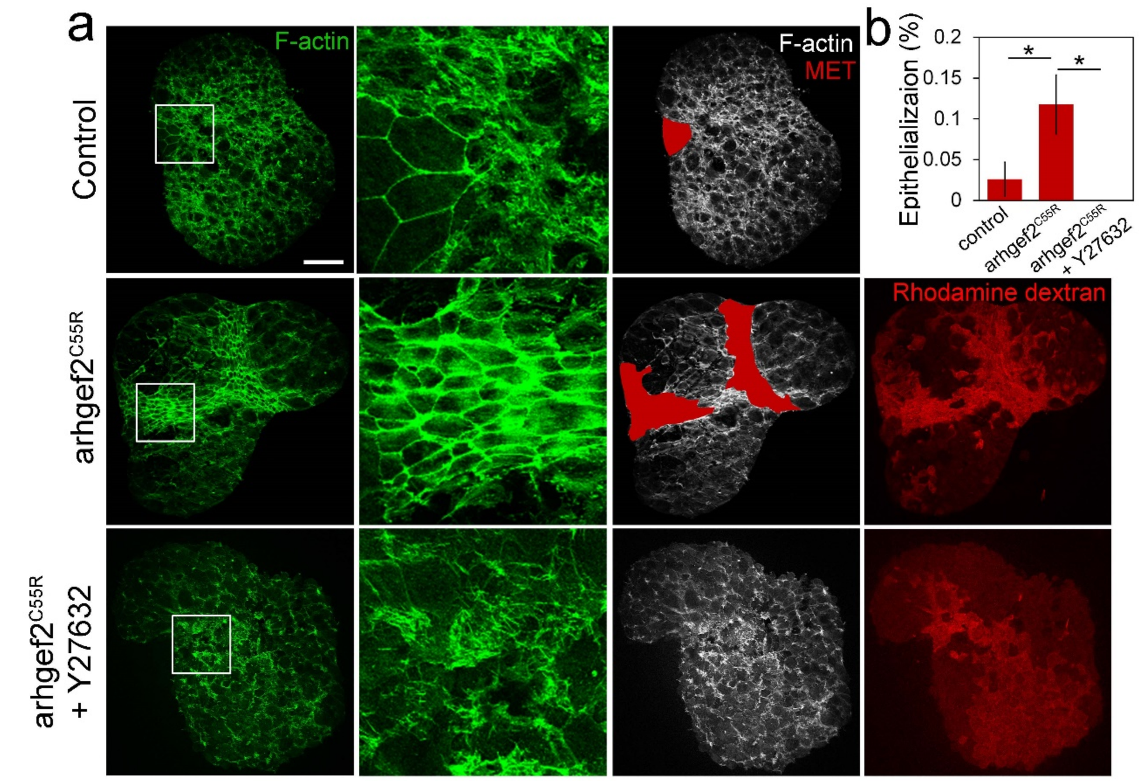


**Supplementary Figure 3: ROCK inhibition blocks epithelialization driven by constitutive active Rho-GEF.**

a. Representative maximum projection confocal images of F-actin labeled aggregates. Epithelialization proceeds in control 5 hpa aggregates is enhanced in arhgef2$^{C55R}$ expressing aggregates. Incubation of arhgef2$^{C55R}$ expressing aggregates in ROCK inhibitor Y27362 inhibits epithelialization.

b. Percent of epithelialization from treatments shown in (a).

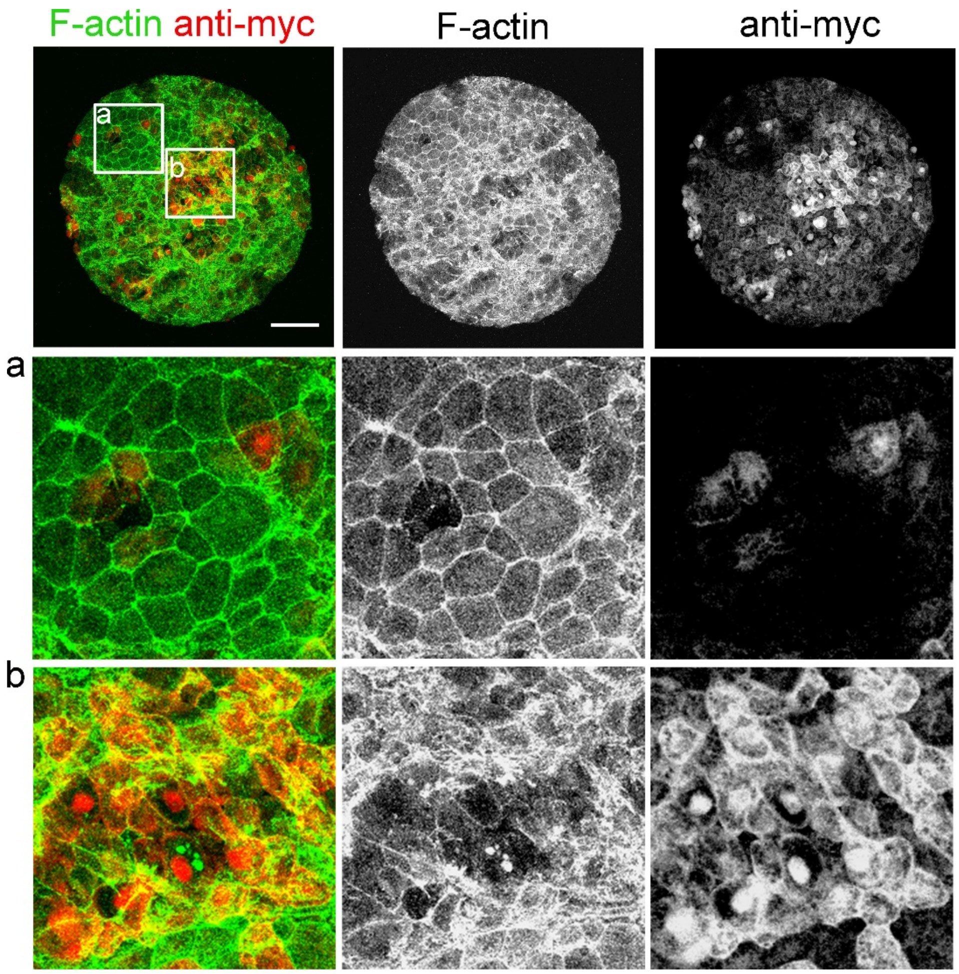


**Supplementary Figure 4: ΔC-C-cadherin expressing cells are not able to transition to epithelial on the surface of deep ectoderm aggregates.** F-actin and myc stained deep ectoderm aggregates at 5 hpa. Cells expressing myc-tagged ΔC-C-cadherin are detected by Anti-myc staining. ΔC-C-cadherin expressing cells (yellow) do not undergo epithelial transition (b) while other cells epithelialize (a) within same aggregate. Scale bar, 100 µm.

### Methods

##### Embryos and embryonic cell aggregates

Embryos were obtained by standard methods; fertilized embryos were cultured in 1/3× modified Barth Solution (MBS; [41]) to the desired stage. To assemble the deep ectoderm aggregates, we first microsurgically isolated ectoderm explants from early gastrula embryos (Stage 10) in Danilchik's For Amy (DFA) medium supplemented with antibiotic and antimycotic (Sigma). Next, ectoderm explants were transferred calcium-magnesium free-DFA which allowed separation of deep and superficial layers of the ectoderm explant. After discarding the superficial layer, freshly isolated deep ectoderm cells were transferred to a PCR tube using a pipette. To provide a non-adherent surface for aggregation, PCR tubes were coated with 1% BSA overnight at 4° C. To confirm that our methods did not produce aggregates contaminated by epithelial cells from the superficial layer, the surface layer of whole embryos was labeled with NHS-Rhodamine (Thermo Scientific)[42] and used to assemble deep ectoderm aggregates (Fig. 1c and d ). Typically, 1 to 2 embryo-ectoderm explants were used to assemble a single aggregate except to make different sizes of aggregates (Fig. 1e). Deep ectoderm aggregates acquire a spherical shape within 2 to 3 hours post aggregation (hpa), at which point they can be subjected to further procedures including treatment with pharmacological inhibitors, measurement of surface tension, and live imaging.

##### Microinjection of localization and overexpression constructs

To visualize or overexpress proteins, mRNAs encoding the desired proteins were transcribed in vitro and injected at the 1- or 2-cell stage of fertilized embryos. GFP-ZO-1 was constructed by moving the entire eGFP-human TJP1 fusion protein obtained from Addgene (Plasmid #30313; Addgene, Cambridge, MA) into pCS2+, mRFP-ZO-1 plasmid provided kindly by Dr. Anne Miller. To induce cell contractility, embryos were injected with arhgef2$^{C55R}$ at ~87.5 pg per embryo [25]. To alter cell-cell adhesion, ΔE-C-cadherin (~1 ng per embryo) and ΔC-C-cadherin (~3.8 ng per embryo) were injected and ectopic gene expression was confirmed by both myc staining and phenotype [21]. Xenopus Yap1.S (formerly YAP1) was amplified from a Stage 10 whole-embryo cDNA library and subcloned C-terminal to eGFP into a pCS2+ vector (InFusion Cloning, Takara, Mountain View, CA).The constitutive active form a myosin binding subunit (MBS T695A) was obtained from the open reading frame of human protein phosphatase 1 regulatory subunit 12A (PPP1R12A, MBS) and amplified from pGEX-PPP1R12A wild type (a gift from Anne Brunet, Addgene plasmid 31669; [43]) and subcloned into pCS2+. The threonine at position 695 was mutated to alanine by PCR sewing using overlapping primers. All constructs were verified by sequencing.

**Histology and immunofluorescence**

Aggregates were fixed with 4% paraformaldehyde and 0.25% glutaraldehyde in PBST (1x PBS with 0.1% Triton X-100) for 15 minutes at room temperature for visualizing F-actin and fibronectin. For immunofluorescence of ZO-1, keratin, aPKC and Itln1, aggregates were fixed with ice cold Dent's fix (4:1 of Methanol: DMSO). Fixed aggregates were washed for 30 minutes with PBST and blocked with 10% goat serum in PBST for 1 hour prior to antibody staining. Primary antibodies for FN (4H2), aPKC (nPKCζ (C-20) sc-216; Santa Cruz), acetylated tubulin (clone 6-11B-1; Sigma), ZO-1 (Invitrogen), keratin (1h5; Developmental Studies Hybridoma Bank), Myc (9E10; Millipore), and Itln1 (gift from Dr. Eamon Dubaissi, 11770-1-AP; proteintech) were used at 1:200 dilution and incubated overnight on a nutator at 4° C. After washing, the samples were incubated with the appropriate secondary antibody at a 1:200 dilution overnight at 4° C. F-actin and cell nuclei were visualized with BODIPY-FL–phallacidin (1:800) and either YO-PRO (1:10000, Invitrogen) or Hoechst 33342 (1:2000, Thermo Fisher), respectively. Cells were evaluated as having undergone epithelialization if they exhibited no F-actin protrusions and had strong circumapical F-actin, e.g. localization for F-actin to cell boundaries, which co-localize with ZO-1 (Supplement Fig. 2).

For YAP nuclear localization studies, aggregates expressing YAP-GFP and ZO-1-RFP were fixed using 4% PFA in PBS for 15 minutes and stained with Hoeschst. For analysis of the images, tiled images were stitched together using an ImageJ plugin [44] and maximum projection images generated. The Hoescht channel was used to segment nuclei using a simple binary threshold. The YAP-GFP channel was used to segment and exclude cells not expressing fluorescent proteins. A custom macro quantified the mean nuclear intensity of YAP-GFP expression of each cell; the cytoplasmic intensity was obtained from an annular region surrounding each nuclei. YAP localization ratios (nuclear to cytoplasmic) are shown in Fig. 3a-c.

**Pharmacological inhibitors**

To alter cell contractility, aggregates at 2 hpa were incubated with Y27632 (50 µM), Blebbistatin (100 µM), or Calyculin A (10 nM) until 5 hpa. Concentrations were chosen from prior studies in Xenopus [24, 25, 45-49]. All cytoskeletal inhibitors were purchased from Calbiochem (Millipore).

**Compliance measurements using micro-aspiration**

To quantify tissue mechanics, aggregates were placed in the high-pressure reservoir of a dual-reservoir micro-aspirator apparatus [19, 50]. Aggregates were gently compressed using hair tools against a 125 μm diameter channel cast across a PDMS block to form a seal across the opening of the channel. A pressure differential at the channel opening was controlled hydrostatically using a computer-controlled syringe pump. After calibrating the channel to obtain zero flow, a small baseline pressure of approximately -3.8 Pa was applied in order to initiate suction and seal the aggregate onto the channel opening. The tissue aggregate was allowed to relax for 5 minutes before the 125-second micro-aspiration protocol was initiated. At the start of the protocol, the aggregate was imaged at 1 frame per second to obtain an initial aspiration length for the tissue. After 5 frames, the syringe pump removed 800 μL of media from the low-pressure reservoir resulting in a -10 Pa suction pressure applied to the aggregate. After 60 seconds of -10 Pa suction, the 800 μ L was media was replaced to return to the baseline pressure and imaging was continued for an additional 60 seconds to confirm the aggregate was properly sealed onto the channel opening. When carrying out repeated measurements on the same sample, the aggregate was rotated to a different position. The tissue boundary was tracked either manually or automatically with Canny-Deriche edge detection [51]. The aspirated length was measured and the data was fit to the power-law model for creep compliance, which has previously been shown to provide an adequate fit for embryonic tissue responses to stress-application [19]. Time-dependent compliance at 30 seconds (fast response) and 60 seconds (steady-state) is reported as the measurement of aggregate mechanical properties. Compliance calculations for each sample are independent of the magnitude of the pressure applied [19]. The suction force of -10 Pa was chosen in order to sufficiently displace aggregate tissues within the resolution of our imaging system without surpassing the critical pressure, thus ensuring measurements are representative of tissue elastic properties [52].

**qPCR**

To evaluate epithelial and mesenchymal gene expression in embryonic tissues, RNA was isolated using the Quick-RNA MiniPrep kit (Zymo Research) with 10 samples each of ectoderm, deep ectoderm aggregates, and superficial ectoderm aggregates at 3, 6 and 24 hours post aggregation. RNA concentrations were measured using the NanoDrop and total RNA was used to create cDNA with the AccuScript High Fidelity 1st Strand cDNA Synthesis

Kit (Agilent Technologies, Inc.). Forward and reverse primers (see table below) were designed for *Xenopus laevis* epithelial genes ZO-1, Cdh1, Itln1 and Krt12, mesenchymal genes FN, VimA and Snail and controls H4A. qPCR protocols were designed and carried out using the Bio-Rad iQ5 thermal cycler and PCR detection system. 10 uL reaction volumes were prepared according to SsoAdvanced Universal SYBR Green Supermix (Bio-Rad) instruction manual. 40 amplification cycles were carried out followed by a melting curve analysis to ensure product purity. Categorization of gene expression at 3 hpa was done using $C_T$ values ($C_T$ > 30 –, 25-30 +, < 25 ++). Fold changes at 24 hpa were calculated using the Livak method, using H4 as the reference gene and 3 hpa as the calibrator [53]. Results are consistent across triplicate experiments with two replicates.

| Gene | Alternate Name | GenBase Name | Accession | Forward Primer | Reverse Primer |
|---|---|---|---|---|---|
| Cdh1 | E-Cadherin | Cdh1.S | XB-GENE-6464305 | GCTGTTGTTGCTCTTACT | CGAGTCTCATCTTCTGGA |
| ZO-1 | TJP-1 | Tjp-1.L | XB-GENE-17332215 | ATATCCAAGCAGTCAGAGA | TCATCTTCATCATCATCTTCC |
| Krt12 | | Krt12.S | XB-GENE-17333623 | TTCCACATCACAATCATCTT | ACCACTTCTTCCACGATA |
| Itln1 | Xeel | Itln1.L | XB-GENE-6256033 | ACTGAGAGGGCTACACTTGCT | ATGTAACCACCTCCTCCAATGC |
| FN | | Fn1.S | XB-GENE-865084 | GGTGGAGGTGTGACAATT | TTGGTATCTCTGTGTAAACTGA |
| VimA | | Vim.L | XB-GENE-866225 | GCTAATCGCAACAATGATG | TTGAATAGTGTCCTGATAGTTAG |
| Snail | | Snail.L | XB-GENE-865328 | GGAGAGTCAGACAGTGTATA | CAAGAGGTGTGTAGTAAGC |
| H4 | | hist1h4a.L | XB-GENE-6493984 | GACGCTGTCACCTACACCGAG | CGCCGAAGCCGTAGAGAGTG |

**Statistical analysis**

Statistical differences of creep compliance among different time points post aggregation were calculated with ANOVA and planned repeated contrast. ANOVA of drug treated aggregates showed data did not vary significantly by clutch ($P > 0.05$), so data was pooled together. Statistical differences between drug treatments were calculated with ANOVA and post hoc analysis. Significance of percentages of epithelialization between control and different experimental conditions were calculated using Mann-Whitney U-test (IBM SPSS, version 22).

40. Chien, Y.-H., Keller, R., Kintner, C. & Shook, D.R. Mechanical Strain Determines the Axis of Planar Polarity in Ciliated Epithelia. *Current Biology* **25**, 2774-2784 (2015).

41. Sive, H.L., Grainger, R.M. & Harland, R.M. (eds.) *Early development of Xenopus laevis: a laboratory manual.* (Cold Spring Harbor Laboratory Press, Cold Spring Harbor, New York; 2000).

42. Edlund, A.F., Davidson, L.A. & Keller, R.E. Cell segregation, mixing, and tissue pattern in the spinal cord of the Xenopus laevis neurula. *Dev Dyn* **242**, 1134-1146 (2013).

43. Banko, M.R. *et al.* Chemical genetic screen for AMPKα2 substrates uncovers a network of proteins involved in mitosis. *Molecular Cell* **44**, 878-892 (2011).

44. Preibisch, S., Saalfeld, S. & Tomancak, P. Globally optimal stitching of tiled 3D microscopic image acquisitions. *Bioinformatics* **25**, 1463-1465 (2009).

45. Zhou, J., Kim, H.Y. & Davidson, L.A. Actomyosin stiffens the vertebrate embryo during critical stages of elongation and neural tube closure. *Development* **136**, 677-688 (2009).

46. Zhou, J., Pal, S., Maiti, S. & Davidson, L.A. Force production and mechanical adaptation during convergent extension. *Development* **142**, 692-701 (2015).

47. Pfister, K., Shook, D.R., Chang, C., Keller, R. & Skoglund, P. Molecular model for force production and transmission during vertebrate gastrulation. *Development* **143**, 715-727 (2016).

48. Rolo, A., Skoglund, P. & Keller, R. Morphogenetic movements driving neural tube closure in Xenopus require myosin IIB. *Dev Biol* **327**, 327-338 (2009).

49. Skoglund, P., Rolo, A., Chen, X., Gumbiner, B.M. & Keller, R. Convergence and extension at gastrulation require a myosin IIB-dependent cortical actin network. *Development* **135**, 2435-2444 (2008).

50. von Dassow, M. & Davidson, L.A. Natural variation in embryo mechanics: gastrulation in Xenopus laevis is highly robust to variation in tissue stiffness. *Dev Dyn* **238**, 2-18 (2009).

51. Schneider, C.A., Rasband, W.S. & Eliceiri, K.W. NIH Image to ImageJ: 25 years of image analysis. *Nat Methods* **9**, 671-675 (2012).

52. Sato, M., Levesque, M.J. & Nerem, R.M. An application of the micropipette technique to the measurement of the mechanical properties of cultured bovine aortic endothelial cells. *Journal of biomechanical engineering* **109(1)**, 27-34 (1987).

53. Šindelka, R., Ferjentsik, Z. & Jonák, J. Developmental expression profiles of Xenopus laevis reference genes. *Developmental Dynamics* **235**, 754-758 (2006).
